## Supplementary Figures for "Systematic evaluation of normalization methods for glycomics data based on performance of network inference"

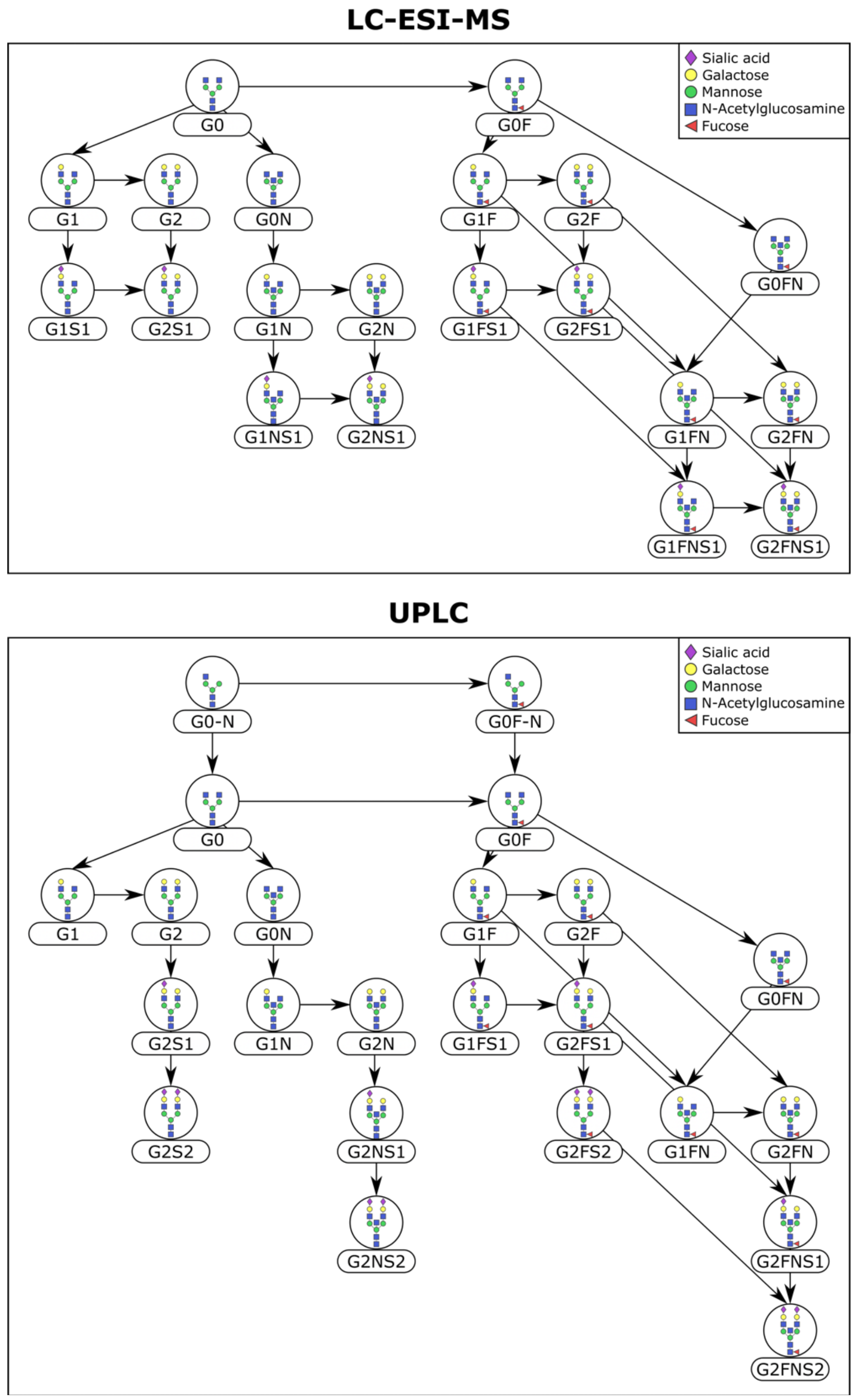


**Figure S1: Reference pathway for IgG LC-ESI-MS and UHPLC-FLD data.** IgG glycans include monosaccharides such as mannose, N-acetylglucosamine, galactose, fucose and sialic acid. Since each of the two platforms allows to measure a slightly different set of IgG glycans, the reference pathway was adapted to fit the structures quantified in the respective dataset.

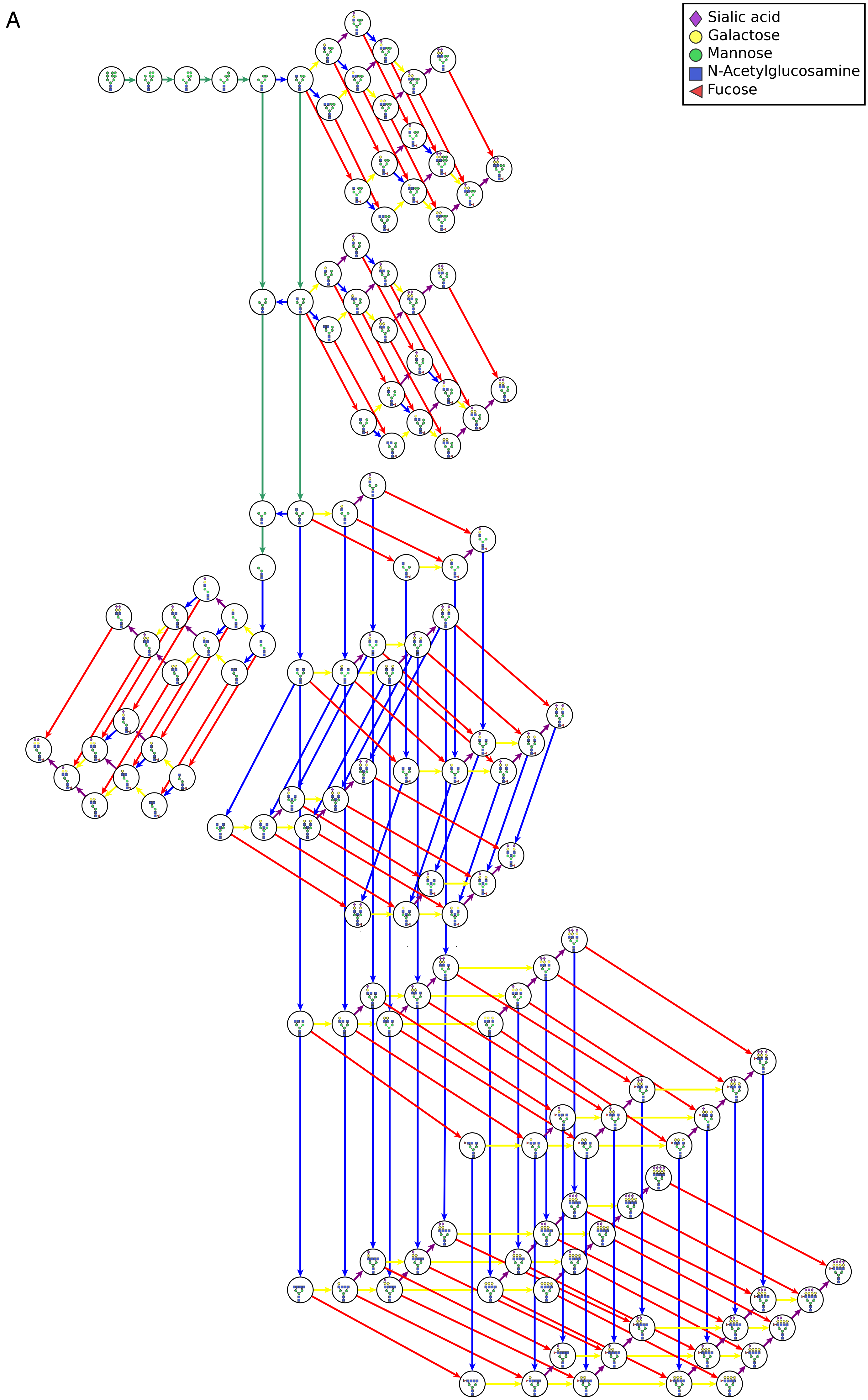
A
Sialic acid
Galactose
Mannose
N-Acetylglucosamine
Fucose

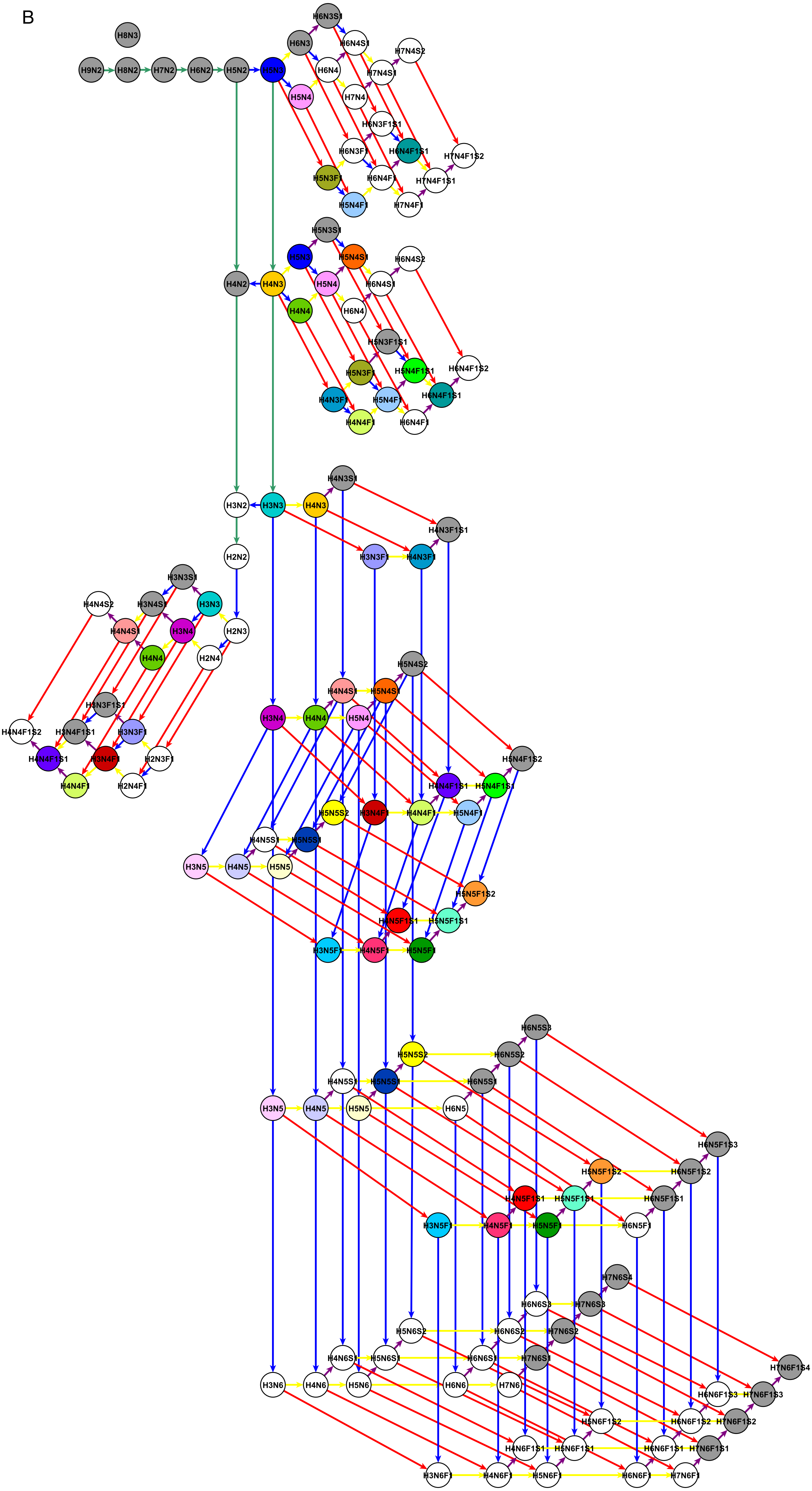
B
H8N3
H9N2
H8N2
H7N2
H6N2
H5N2
H5N3
H6N3S1
H6N3
H6N4S1
H7N4S2
H6N4
H7N4S1
H5N4
H7N4
H6N3F1S1
H6N3F1
H6N4F1S1
H7N4F1S2
H5N3F1
H6N4F1
H7N4F1S1
H5N4F1
H7N4F1
H5N3S1
H5N3
H5N4S1
H6N4S2
H4N2
H4N3
H5N4
H6N4S1
H4N4
H6N4
H5N3F1S1
H5N3F1
H5N4F1S1
H6N4F1S2
H4N3F1
H5N4F1
H6N4F1S1
H4N4F1
H6N4F1
H4N3S1
H3N2
H3N3
H4N3
H4N3F1S1
H2N2
H3N3F1
H4N3F1
H3N3S1
H4N4S2
H3N4S1
H3N3
H4N4S1
H3N4
H2N3
H4N4
H2N4
H3N3F1S1
H4N4F1S2
H3N4F1S1
H3N3F1
H4N4F1S1
H3N4F1
H2N3F1
H4N4F1
H2N4F1
H5N4S2
H4N4S1
H5N4S1
H3N4
H4N4
H5N4
H5N4F1S2
H4N4F1S1
H5N4F1S1
H5N5S2
H3N4F1
H4N4F1
H5N4F1
H4N5S1
H5N5S1
H3N5
H4N5
H5N5
H5N5F1S2
H4N5F1S1
H5N5F1S1
H3N5F1
H4N5F1
H5N5F1
H6N5S3
H5N5S2
H6N5S2
H4N5S1
H5N5S1
H6N5S1
H3N5
H4N5
H5N5
H6N5
H6N5F1S3
H5N5F1S2
H6N5F1S2
H4N5F1S1
H5N5F1S1
H6N5F1S1
H3N5F1
H4N5F1
H5N5F1
H6N5F1
H7N6S4
H6N6S3
H7N6S3
H5N6S2
H6N6S2
H7N6S2
H4N6S1
H5N6S1
H6N6S1
H7N6S1
H7N6F1S4
H3N6
H4N6
H5N6
H6N6
H7N6
H6N6F1S3
H7N6F1S3
H5N6F1S2
H6N6F1S2
H7N6F1S2
H4N6F1S1
H5N6F1S1
H6N6F1S1
H7N6F1S1
H3N6F1
H4N6F1
H5N6F1
H6N6F1
H7N6F1

**Figure S2: Hypothetical pathway of total plasma N-glycan synthesis.** Each node corresponds to a glycan structure. Edges represent possible enzymatic reactions of glycan synthesis, namely all single monosaccharide modifications against which there is no strong experimental evidence against. **A** In each node, the corresponding glycan structure is depicted. **B** Each node is labelled with the composition of the corresponding structure. White nodes indicate that the corresponding composition was not included in the dataset. Gray nodes indicate compositions that can be attributed to a single glycan structure. Colored nodes indicate that there are multiple structures with the same composition.

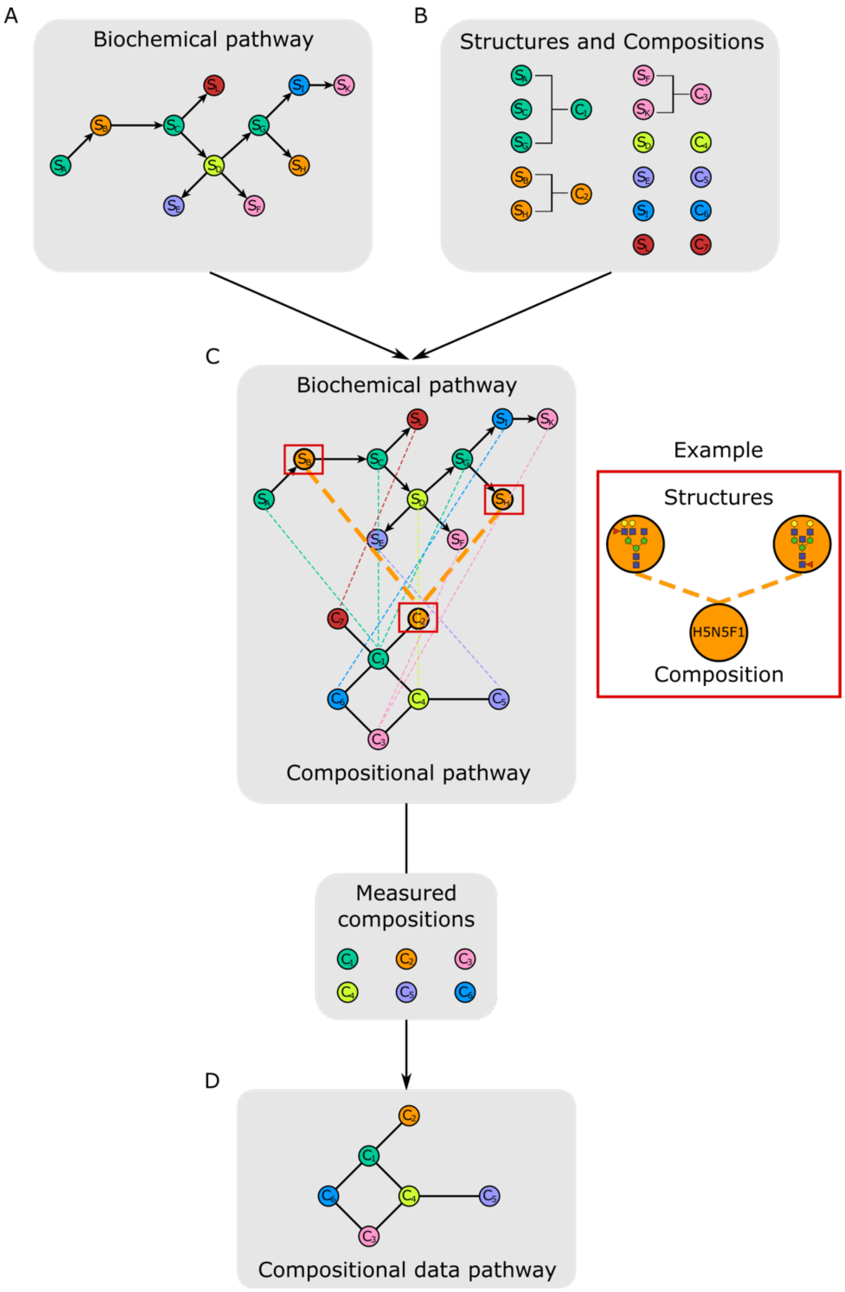


**Figure S3: Creation of the reference pathway for the MALDI-FTICR-MS data. A** Hypothetical biochemical pathway. **B** Mapping between structures in the hypothetical pathway and the corresponding compositional name. **C** Structures with the same mass in the canonical pathway were merged into one single compositional node. **D** Since not all compositions in the canonical pathway were measured in the available dataset, the unmeasured compositions were removed from the pathway together with all their edges.

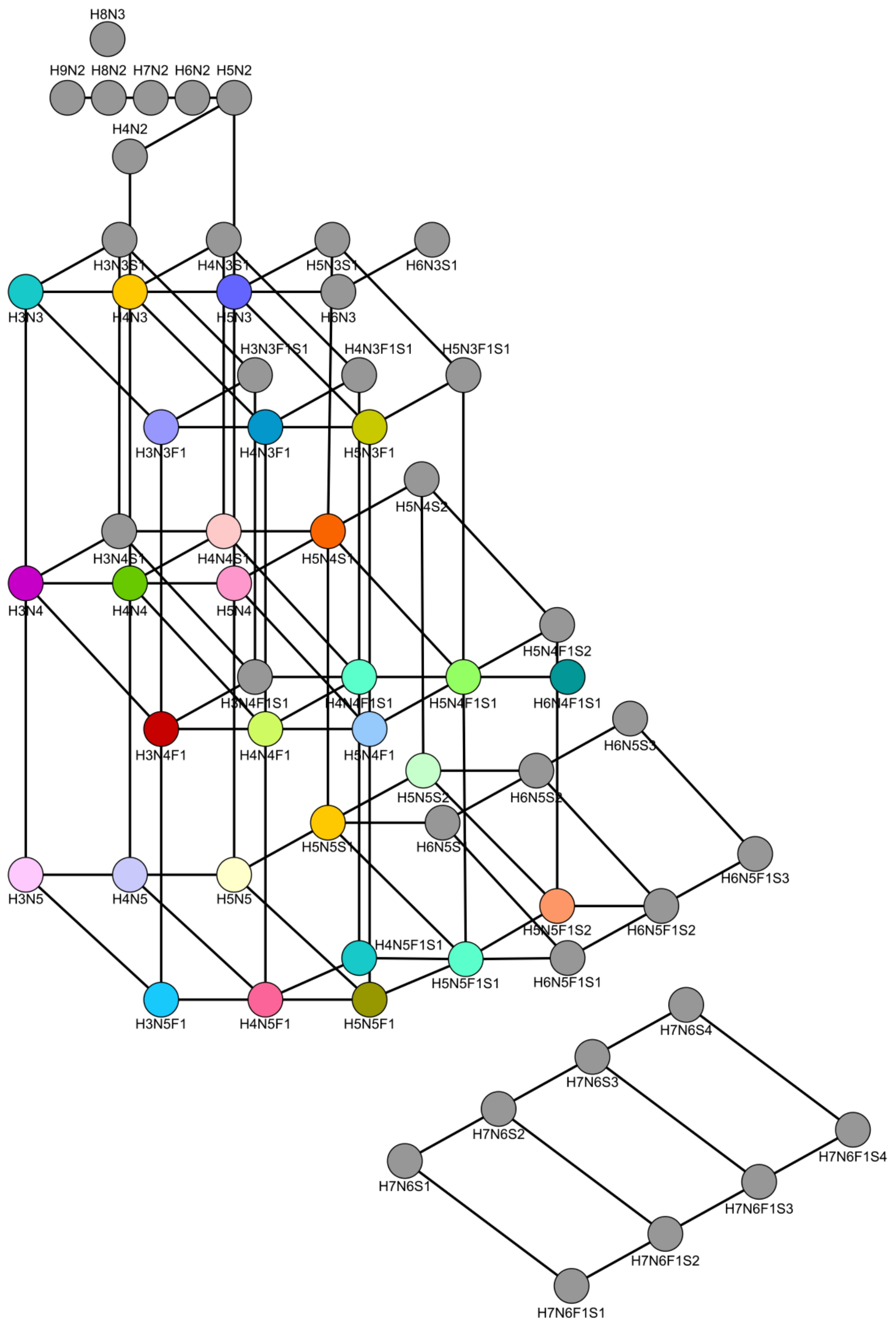


**Figure S4: Reference pathway for total plasma N-glycans in MALDI-FTICR-MS data.** This technology separates glycans based on their total mass, which can then be characterized according to the contained number of hexoses (like glucose, galactose and mannose, indicated with H), N-acetylhexosamines (N-acetylglucosamine, indicated with N), fucoses (F), and sialic acids (S). Grey nodes represent compositions made of a single structure, while colored nodes represent compositions made of more than one structure, as indicated in Figure S2B.

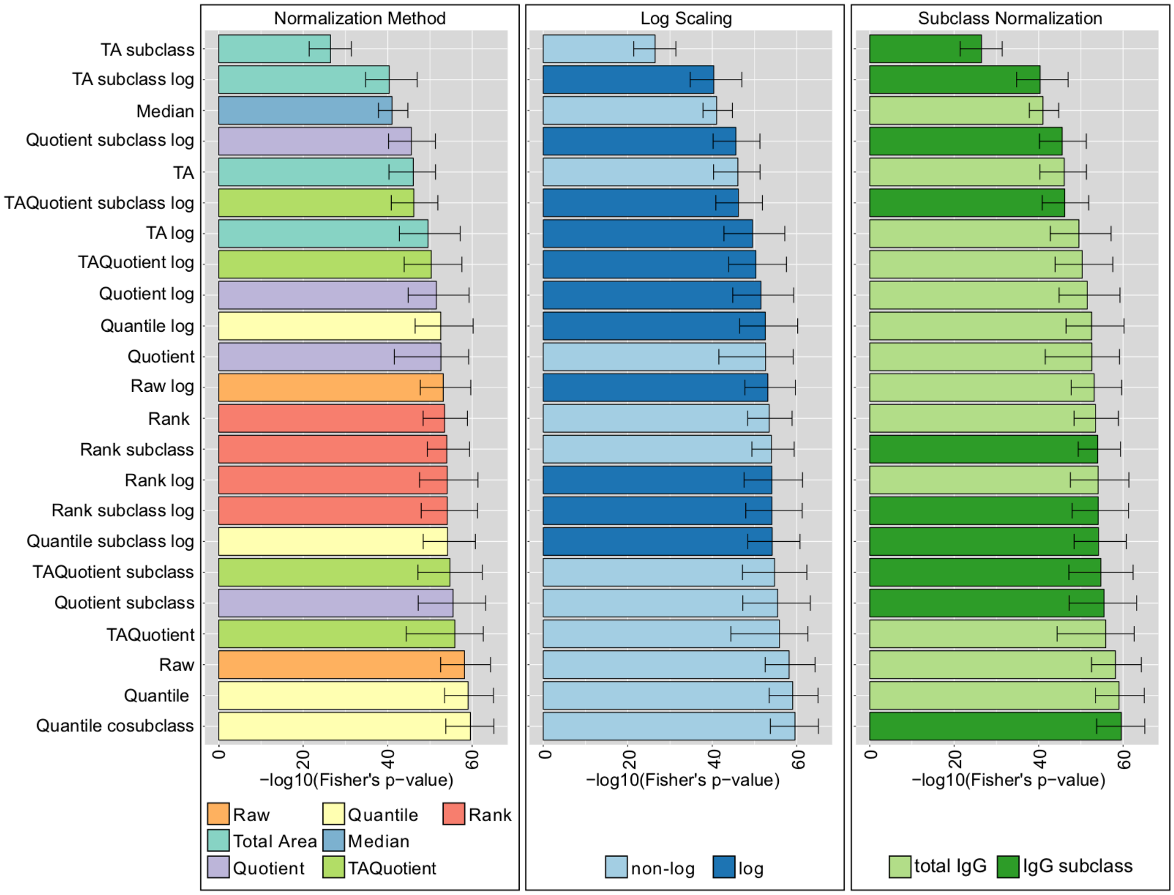


**Figure S5: LC-ESI-MS normalization analysis results (Korčula 2010 cohort).** Results in the panels are colored according to type of normalization (left), log-transformation (center), or normalization per IgG subclass or total IgG (right). Bars represent the median of the Fisher's exact test p-values over 1,000 bootstrapping, and error bars the corresponding 95% confidence intervals.

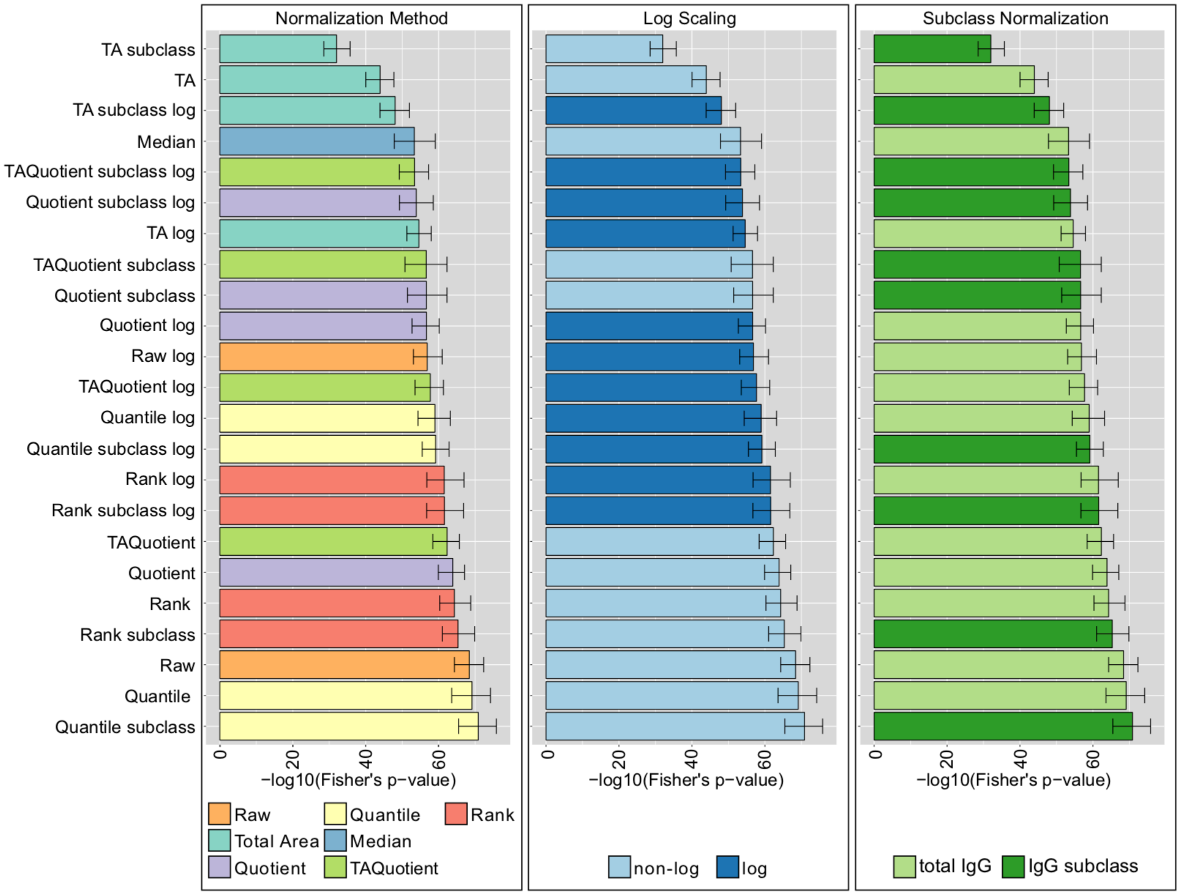


**Figure S6: LC-ESI-MS normalization analysis results (Split cohort).** Results in the panels are colored according to type of normalization (left), log-transformation (center), or normalization per IgG subclass or total IgG (right). Bars represent the median of the Fisher's exact test p-values over 1,000 bootstrapping, and error bars the corresponding 95% confidence intervals.

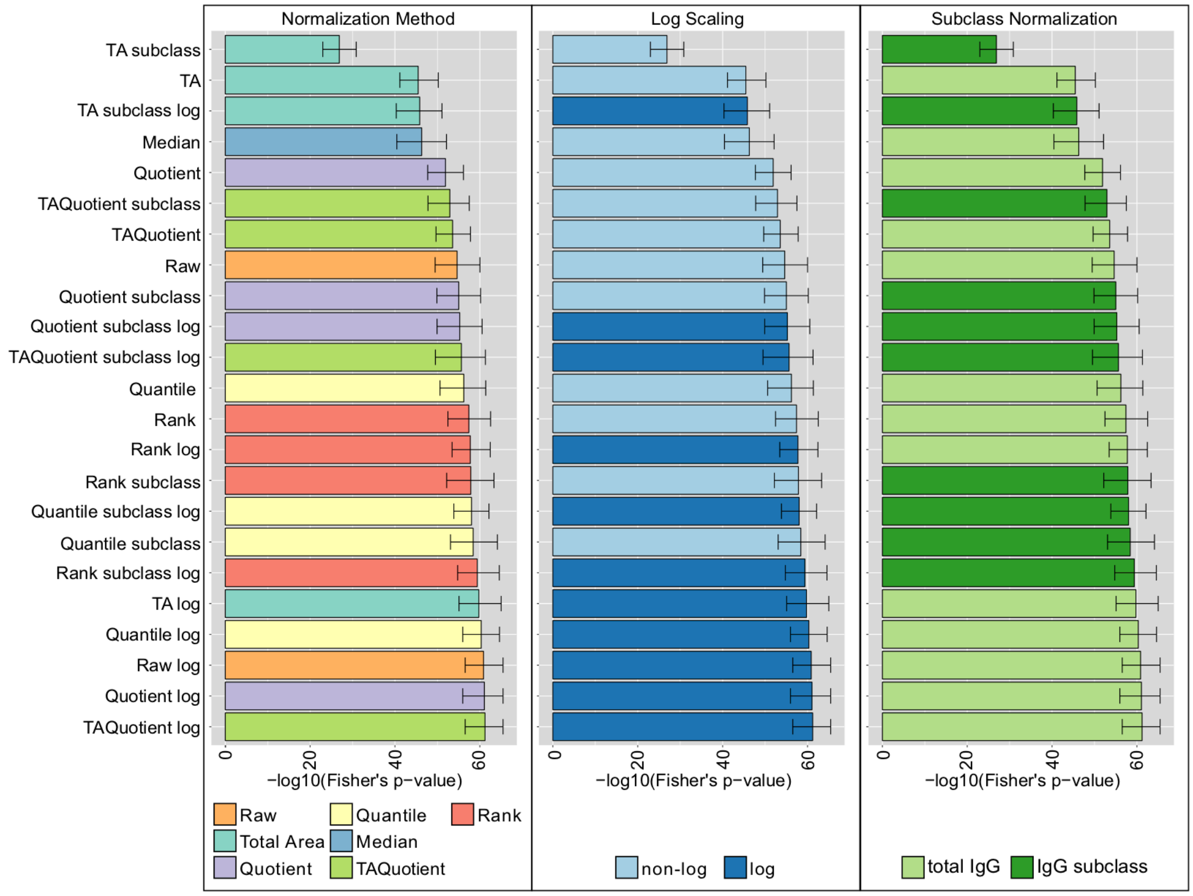


**Figure S7: LC-ESI-MS normalization analysis results (Vis cohort).** Results in the panels are colored according to type of normalization (left), log-transformation (center), or normalization per IgG subclass or total IgG (right). Bars represent the median of the Fisher's exact test p-values over 1,000 bootstrapping, and error bars the corresponding 95% confidence intervals.
